# Effects of treadmill exercise on retinal vascular morphology, function, and circulating immune factors in a mouse model of retinal degeneration

**DOI:** 10.1101/2025.02.11.637694

**Authors:** Hayden B. Haupt, John M. Nickerson, Jeffrey H. Boatright, Machelle T. Pardue, Katie L. Bales

## Abstract

**Purpose:** Exercise is neuroprotective in rodents undergoing retinal degeneration (RD). However, the effects of exercise on retinal vasculature remain unexplored. Here, we investigate whether treadmill exercise influences retinal vascular morphology, function, gene expression, and circulating factors in a light-induced retinal degeneration (LIRD) mouse model.

**Methods:** 6-week-old female BALB/c mice were assigned to inactive+dim, active+dim, inactive+LIRD and active+LIRD groups (n=20 per group). Active mice were treadmill exercised (1hr/d 10m/min) for two weeks, then LIRD was induced (5000 lux/4hrs). Inactive mice were placed on a static treadmill. Retinal neurovascular coupling was measured with functional hyperemia (FH) and vascular morphology using OCT-A. Vascular gene expression was quantified from isolated retinal endothelial cells using ddPCR five days following LIRD. Serum was collected for circulating cytokine and chemokine analyses. Data were analyzed using 2-way ANOVA.

**Results:** Retinal vessel vasodilation was significantly increased in active+LIRD mice compared to inactive+LIRD mice. Superficial and intermediate/deep vascular plexi from inactive+LIRD mice had significantly decreased vessel density and total vessel length, with increased numbers of end points and lacunarity compared to active groups. Isolated retinal endothelial cell gene expression varied among groups. Most notably, Active+LIRD mice had a distinct immune response profile, with increased expression of IL-6, KC, and VEGF-A.

**Conclusions:** Treadmill exercise maintained retinal vascular morphology and function, modestly altered endothelial gene expression, and is associated with a specific circulating immune response profile in a LIRD mouse model. These data indicate therapeutic effects of exercise on retinal vasculature in RD.

## Introduction

The structure of the retinal vasculature is optimally designed to meet the constant metabolic demands of retinal neurons. The retina has two main sources of blood supply, both of which originate from the ophthalmic artery: the choroidal blood vessels which supply photoreceptors whereas the central retinal artery supports the inner retina^1^. Several studies have correlated reduced blood flow and or atrophy of the retinal capillary network with photoreceptor loss in animal models and in patients with retinal degenerative diseases^2,3^. In all cases, this reduced retinal capillary blood flow was associated with significant reductions in vascular diameter and the presence of nonperfused blood vessels^3^. Within RD, vascular deficits have been described as an aftereffect of retinal cell dysfunction and have been associated with late-stage disease^4,5^. Emerging research suggests that retinal vascular deficits occur earlier in RD pathogenesis^2,6^. These studies confirm that vascular dysfunction in retinal degeneration is not a late-stage consequence of neuronal dysfunction but is present early in disease development.

Our previous work has thoroughly demonstrated exercise as an effective neuroprotective method in several animal models of RD, preserving photoreceptor function, RPE integrity, and retinal astrocyte morphology^7–11^. However, few studies have evaluated retinal vasculature in models of photoreceptor degeneration with exercise interventions and none have measured vascular function^12^. Accordingly, the current study investigates whether treadmill exercise influences retinal vascular morphology, function, endothelial gene expression and circulating factors in healthy and degenerating retinas.

## MATERIALS AND METHODS

### Animals

All animal procedures were approved by the Atlanta VA Institutional Animal Care and Use Committee and conform to the ARVO Statement for the Use of Animals in Ophthalmic and Vision Research. Adult BALB/c female mice were purchased from Charles River (8–10 weeks old; Wilmington, MA, USA) and housed under a 12:12 light: dark cycle with ad libitum access to water and standard mouse chow (Teklad Global 18% Protein Rodent Diet 2918, Irradiated, Rockville, MD).

### Experimental design

Mice were randomly assigned to one of the following four groups: inactive+dim, active+dim, inactive+LIRD, and active+LIRD (n=20 per group). Active groups ran on a rodent treadmill once daily at 10 m per minute (m/min), 5 days per week for 3 weeks (**Figure 1A**). Inactive groups were placed on a static treadmill for the same amount of time. On the day of LIRD, mice were exposed to toxic light within 30 min after treadmill activity. Following 1 additional week of treadmill running, functional hyperemia (FH) and optical coherence tomography angiography (OCT-A) was performed to assess retinal vascular function and morphology *in vivo*, respectively. Mice were euthanized via CO_2_ gas inhalation and secondary cervical dislocation, retinas were collected and pooled together by cohort for CD31+ cell isolation via magnetic activated cell sorting (MACS). Serum was isolated from whole blood. Samples were collected and stored at −80 degrees.

**Figure 1.**
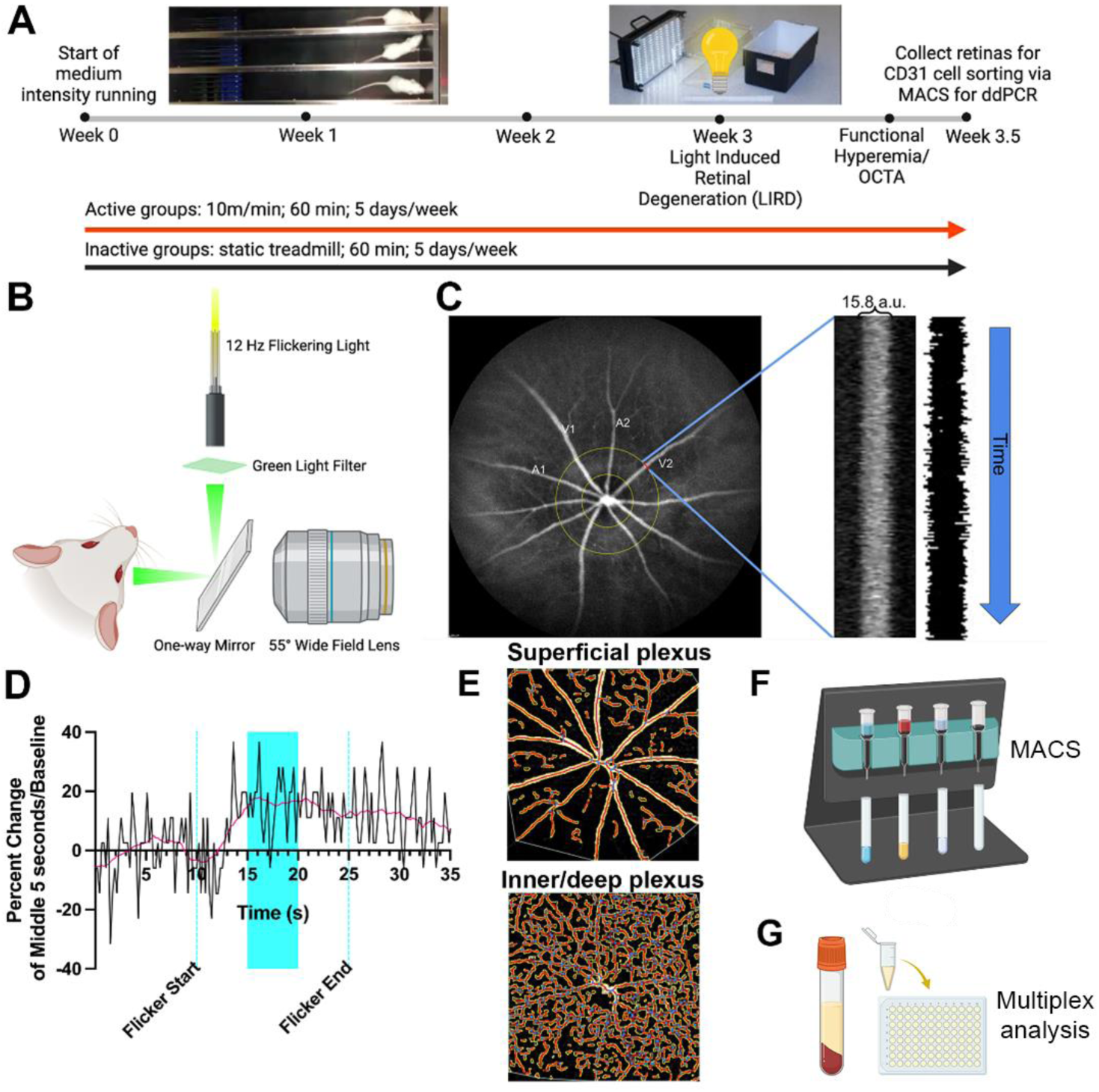
Experimental methods and timeline. 6-week-old female BALB/c mice were assigned to inactive+dim, active+dim, inactive+LIRD and active+LIRD groups (n=20 per group). Active mice were treadmill exercised (1hr/d, 10m/min) for two weeks, then LIRD was induced (5000 lux/4hrs). Inactive mice were placed on a static treadmill for the same schedule (**A**). Retinal neurovascular coupling was measured by functional hyperemia. Eyes were illuminated using a 12 Hz flickering green light (**B**). Scanning laser ophthalmoscope image identifying whether a vessel is a venule (Vx) or arteriole (Ax), (**C**). Plot of vessel diameter across time used to measure the percent change of vessel caliber during stimulation (highlighted region; **D**). OCT-A images showing vascular morphology quantification following Angiotool analysis (**E**). 5-days following LIRD, retinal extracts were pooled to perform (**F**) magnetic-activated cell sorting (MACS) to isolate retinal endothelial cells. ddPCR was performed to quantify vascular gene expression. Serum from all experimental groups was analyzed via multiplex assay (**G**).

### Exercise regimen and light exposure

In accordance with previous studies, active mice ran 60 min per day between ZT3-5 on treadmills equipped with electric shock gratings (Exer-3/6; Columbus Instruments, Columbus, OH, USA)^9,10^. Inactive mice were placed on static treadmills. Following 2 weeks of exercise, the mice were exposed to typical laboratory lighting (50 lux; dim) or toxic light (5000 lux; LIRD) for 4h using a light emitting diode (LED) light panel (LED500A; Fancierstudio, Hayward, CA, USA). This level of toxic light is a moderate brightness to induce retinal degeneration. For light exposure, animals were individually housed in shoebox containers with the LED light panel placed above, as previously described^9,10^. Room and light box temperatures were closely monitored to ensure animal welfare.

### Functional hyperemia

To induce a maximal functional hyperemia response, filtered light (480-600 nm) from a fiber optic illuminator was gated with an electromechanical shutter (Optical Beam Shutter; Thorlabs, Newton, NJ) at 12 Hz (**Figure 1B**). The flicker light was reflected off a 45-degree angle prism mirror (TS Cold Mirror; Edmundoptics, Barrington, NJ) focused onto the eye, with a luminance of 5000 lux at the surface of the eye. Animals were anesthetized via an intraperitoneal injection (ketamine [80 mg/kg]/xylazine [16 mg/kg]) and retinal blood vessels visualized via an intraperitoneal injection of indocyanine green (ICG) dye (ICG, 20 mg/kg; IC-Green; Akorn, Lake Forest, IL). The pupils were dilated (1% tropicamide; Alcon Laboratories, Ft. Worth, TX, USA) and corneas were anesthetized (1% tetracaine; Alcon Laboratories, Ft. Worth, TX, USA). Finally, contact lenses were placed on the cornea with Systane (Alcon, Ft. Worth TX, USA) eye drops to keep the eyes hydrated during imaging. Retinal blood vessels were continuously monitored with a scanning laser ophthalmoscope using a 55° wide field lens (SLO; Heidelberg Spectralis; (Heidelberg Engineering, Heidelberg Germany) during flicker stimulation. Each stimulation trial consisted of 10 sec of baseline measurement (without flicker), followed by 15 sec of stimulation with flicker light, and ending with 10 sec of recovery assessment.

Measurement and quantification of the flicker-induced vasodilation was performed on ICG fundus images using specialized imaging software (ImageJ; NIH, Bethesda, MA). A line was drawn perpendicular to a first-order arteriole or venule at one optic disk distance away from the optic nerve. Then, the ICG-filled lumen of the vessel that intersected with the line was extracted over the entire stimulation trial, generating a distance (vessel caliber) vs. time line scan image (also known as kymograph). To calculate the percent vasodilation, we averaged the vessel diameters at 0-10 sec (baseline) and 10-25 sec (stimulation) and measured the percent change in vessel caliber from baseline to stimulation (**Figure 1C,D**). Calculations are performed on two arterioles or venules per retina for each animal at each time-point before averaging. Resting vessel caliber was measured one optic nerve diameter from the disk along the vessel length and normalized to the optic disk diameter to eliminate any magnification error. The recovery phase was not included in the measurement but served to show the success of flicker-induced vasomotor response. Plots of the functional hyperemic response were smoothed with a moving window average smoothing algorithm (Graphpad Prism 10.2.3, Boston, MA, USA (**Figure 1D**).

### Optical Coherence Tomography Angiography (OCT-A)

Optical Coherence Tomography Angiography (OCT-A) was performed in conjunction with the functional hyperemia session. Anesthetized animals were imaged using a scanning laser ophthalmoscope with a 30° animal lens (SLO; Heidelberg Spectralis; (Heidelberg Engineering, Heidelberg Germany). Images were obtained with the high-speed setting at 30° IR, 20° scan angle, 20° x 20° scan area and consisted of 512 B-scans at 7µm increments. Images below a quality threshold of 28 as determined by the Heidelberg Spectralis were excluded from analysis. OCT-A images of the default superficial vascular complex (SVC) and the intermediate/deep vascular complexes (I/DVC) were delineated automatically and were used for analyses. Images with movement caused by heavy breathing or other artifacts were excluded from analysis. Images were exported as 8-bit images and then processed using Angiotool for vascular morphological quantifications with the following parameters: blood vessel diameter (2-25μm) and pixel intensity (0-255; **Figure 1E**)^13^.

### Magnetic Activated Cell Sorting (MACS)

Retinal endothelial cell isolation was performed as previously described^5^. Retinal extract from mice in the same experimental group were pooled together, microdissected, and placed in ice-cold physiological solution (artificial cerebral spinal fluid, aCSF) containing: 125 mM NaCl, 3 mM KCl, 1 mM MgCl_2_, 0.2 mM CaCl_2_, 1.25 mM NaH_2_PO_4_, 25 mM NaHCO_3_, 25 mM glucose and saturated with carbogen (95% O_2_-5% CO_2_ mixture; pH 7.4)^10,14^. Retinas were minced and enzymatically dissociated with a Papain Dissociation kit (Worthington, Lakewood, NJ, USA) following manufacturer’s instructions. Retinal endothelial cells were then isolated using anti-CD31+ (PECAM1; platelet and endothelial cell adhesion molecule 1) MicroBead kit (Miltenyi Biotec, Cambridge, MA, USA). Manufacturer instructions were generally followed, with the exception that incubation times were extended to 15 min and the total volume of microbeads was increased to 35 μl (**Figure 1F**).

### Isolated retinal endothelial cell gene expression

Isolated retinal endothelial cells were probed to measure gene expression associated with angiogenesis and neuroprotection. Isolated cells were flash frozen and RNA was extracted using the Qiagen RNAeasy Mini Kit (Cat. No. 74104, Qiagen LLC, Germantown, MD, USA). QuantiNova cDNA synthesis kit (Cat. No. 205413, Qiagen LLC, Germantown, MD, USA) was used to make cDNA as per the manufacturer’s protocol. Digital droplet polymerase chain reaction (ddPCR) was used to determine relative quantities of transcripts for the genes of interest. ddPCR was performed using 5 ng of cDNA and fluorescent amidite matrix (FAM) hydrolysis probe sets for Vascular Endothelial Growth Factor Receptor 1 (VEGFR1), VEGFR2, Vascular Cell Adhesion Molecule 1 (VCAM1), Endothelin 1 (EDN1), Nitric Oxide Synthase 3 (NOS3), Brain Derived Neurotrophic Factor (BDNF), Nuclear Factor Kappa B Subunit 1 (NFKB1), and Nuclear Factor Kappa B Subunit 2 (NFKB2) and a hexachloro-fluorescein (HEX)-labeled probe assay for hypoxanthine-guanine phosphoribosyltransferase (HPRT) (Integrated DNA Technologies [IDT], Coralville, IA, USA) (**Table 1**). Data were analyzed using QuantaSoft analysis software (Bio-Rad, Hercules, CA, USA), which uses a Poisson distribution model to calculate the number of starting target template molecules in each well from the number of FAM- and HEX-positive droplets.

**Table 1.**
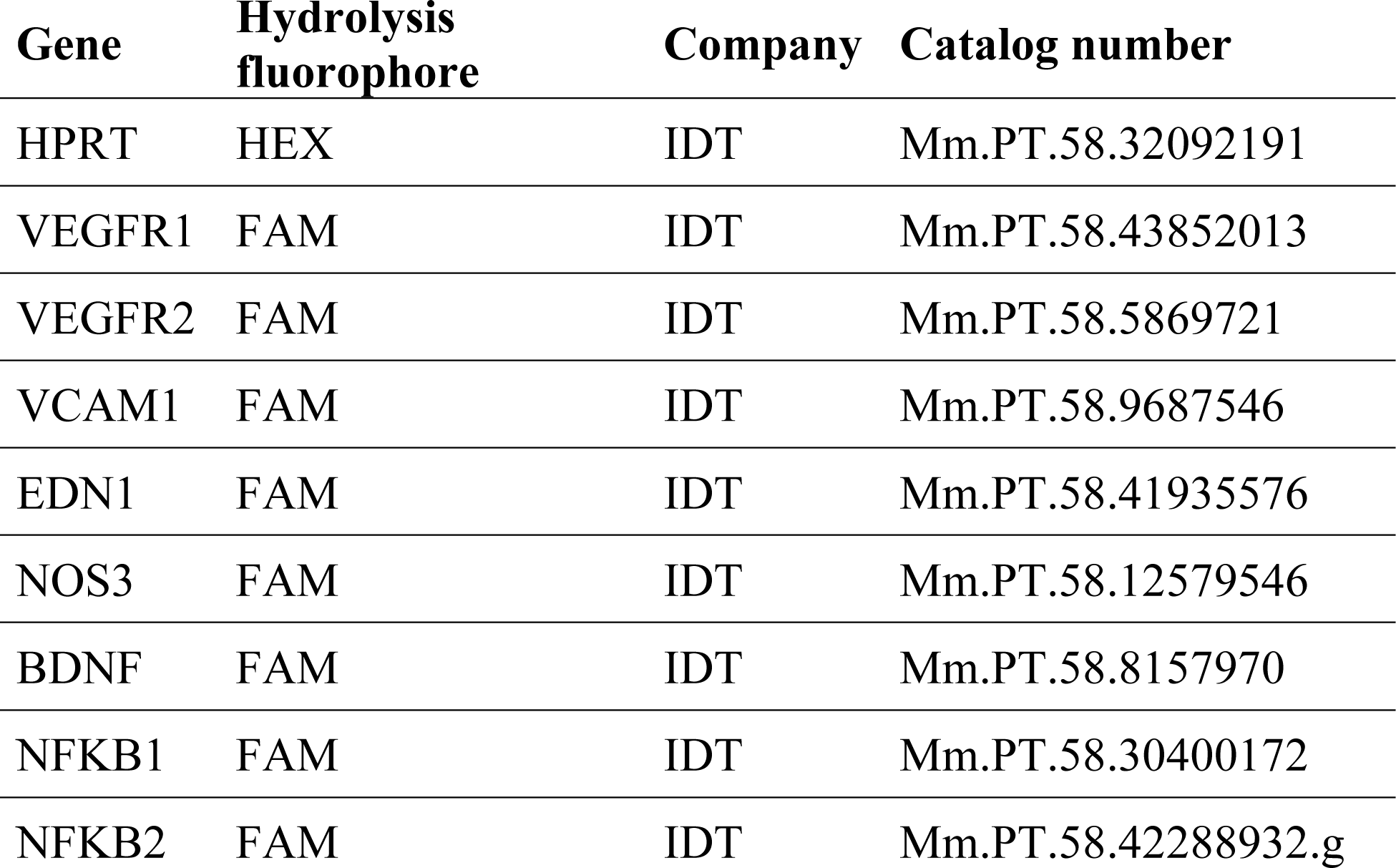
ddPCR Probes.

| Gene | Hydrolysis fluorophore | Company | Catalog number |
| --- | --- | --- | --- |
| HPRT | HEX | IDT | Mm.PT.58.32092191 |
| VEGFR1 | FAM | IDT | Mm.PT.58.43852013 |
| VEGFR2 | FAM | IDT | Mm.PT.58.5869721 |
| VCAM1 | FAM | IDT | Mm.PT.58.9687546 |
| EDN1 | FAM | IDT | Mm.PT.58.41935576 |
| NOS3 | FAM | IDT | Mm.PT.58.12579546 |
| BDNF | FAM | IDT | Mm.PT.58.8157970 |
| NFKB1 | FAM | IDT | Mm.PT.58.30400172 |
| NFKB2 | FAM | IDT | Mm.PT.58.42288932.g |

### Multiplex assay

Serum was collected and analyzed using the U-PLEX Biomarker Group 1 (mouse) 50-Plex (Meso Scale Delivery, Rockville MD, USA) to assess serum cytokine and chemokine expression related to immune response and regulation. Samples were prepared and analyzed per the manufacturer instructions. The following chemokines and cytokines were analyzed: 6CKine/CCL21, BAFF*, BCA-1/BLC, CD40/TNFRSF5, Eotaxin, EPO, GM-CSF, IFN-α, IFN-β, IFN-γ, IL-1β, IL-2, IL-4*, IL-5, IL-6, IL-9*, IL-10, IL-12/IL-23p40, IL-12p70*, IL-13*, IL-15*, IL-16, IL-17A, IL-17A/F, IL-17C, IL-17E/IL-25, IL-17F*, IL-21, IL-22, IL-23*, IL-27p28/IL-30*, IL-31*, IL-33, IP-10, KC/GRO, MCP-1, MCP-5/CCL12, MDC, MIP-1α*, MIP-1β, MIP-2, MIP-3α, MMP-9 (total), NGAL/LCN2*, RANTES, SDF-1α, TARC, TNF-α, TNF-RI, VEGF-A. Chemokines and cytokines with an asterisk were not detectable (**Figure 1G**).

### Masking and Statistical analysis

Sample size was determined based on our previously reported data^8–10^. Researchers who analyzed the data were masked to the experimental procedures and specific treatment groups. For FH, three graders were trained to identify images of acceptable quality, as well as two clearly viewed arteries and venules. If two out of the three graders agreed on the same arteries and venules, they would be analyzed. All data are presented as mean ± standard error of the mean (SEM). Statistical analyses were performed using Graphpad Prism 10.2.3 (San Diego, CA, USA). Two-way ANOVAs on main effects and interactions of exercise and light exposure were performed with Tukey’s multiple comparison tests. All *p*-values lower than .05 were considered statistically significant. The ROUT method (with Q set to 1%) was used to detect outliers.

## RESULTS

### Retinal vascular morphology is preserved in exercised mice undergoing light induced retinal degeneration

Within the superficial vascular plexus (**Figure 2A-H**), inactive+LIRD mice had a significant decrease in vessel density and total vessel length compared to active+dim groups, whereas no statistical differences were observed comparing active+LIRD amongst all groups (vessel density: **Figure 2E**, inactive+dim: 18.98%±0.74; active+dim: 22.02%±0.81; inactive+LIRD: 16.36%±0.98; active+LIRD: 18.62%±1.43; two-way ANOVA, effect of exercise*LIRD, F_(3,42)_=5.58, p=0.0026; total vessel length: **Figure 2F**, inactive+dim: 3842μm ±215.40; active+dim: 4486μm ±175.70; inactive+LIRD: 3450μm±195.30; active+LIRD: 3888μm ±293.20; two-way ANOVA, effect of exercise*LIRD, F_(3,42)_=4.54, p=0.0076). Inactive+LIRD retinas also had increased total number of end points and lacunarity, or regions that do not contain vasculature, while the active+LIRD retinas were protected from these changes (total end points: **Figure 2G**, inactive+dim: 146.90a.u.±8.79; active+dim: 149.40a.u.±5.59; inactive+LIRD: 190.80a.u.±15.85; active+LIRD: 144.50a.u.±10.48; two-way ANOVA, effect of exercise*LIRD, F_(3,42)_=4.54, p=0.0076; lacunarity: **Figure 2H**, inactive+dim: 0.188a.u.±0.0083; active+dim: 0.187a.u.±0.013; inactive+LIRD: 0.266a.u.±0.023; active+LIRD: 0.18a.u.±0.014; two-way ANOVA, effect of exercise*LIRD, effect of exercise*LIRD, F_(3,42)_=8.97, p=0.0001).

**Figure 2.**
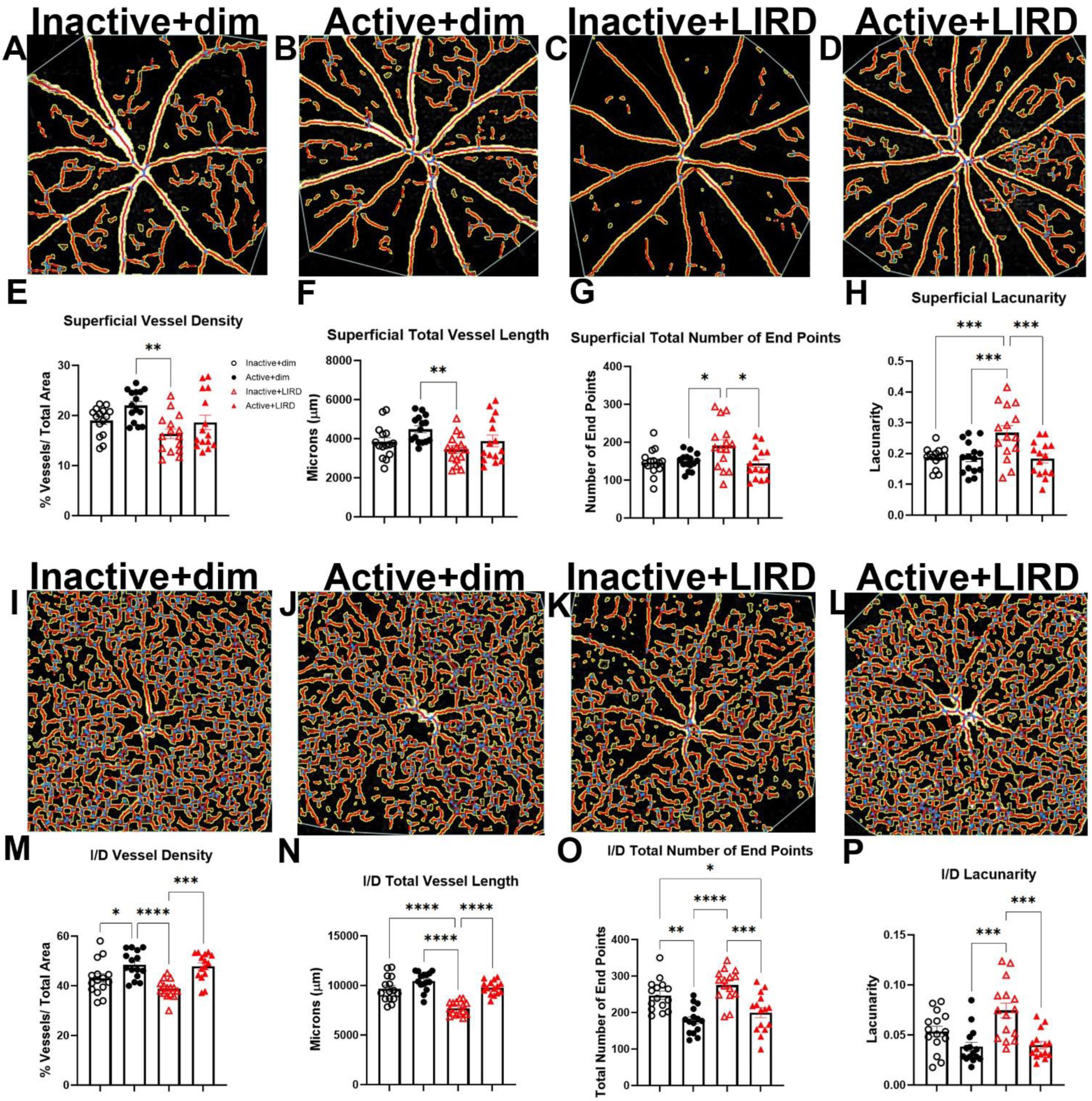
OCT-A reveals Active+LIRD mice have maintained vascular plexi morphology. Optical Coherence Tomography Angiography (OCT-A) was performed to evaluate superficial (**A-D**) and intermediate/ deep (**I-L**) retinal vascular plexi morphology in vivo. Inactive+LIRD mice displayed decreases in vessel density (**E,M**) and total vessel length (**F,N**). However, they had an increase in the total number of end points (**G,O**) compared to active animals in the superficial vascular plexus and in the intermediate/deep. Inactive+LIRD mice had a significant increase in lacunarity (gaps, or regions without vasculature; **H,P**) in both the superficial and intermediate/deep vascular plexi. Two-way ANOVA with Tukey’s multiple comparison analyses were performed, n=15, *p<0.05, **p<0.01, ***p<0.001, p<0.0001.

Intermediate/deep vascular plexi (**Figure 2I-P**) quantifications revealed similar trends as described in the superficial vascular plexus, with inactive+LIRD mice showing greater changes and active+LIRD mice remaining statistically similar to inactive+dim animals. Inactive+LIRD retinas had a significant decrease in vessel density (**Figure 2M**, inactive+dim: 43.12%±1.76; active+dim: 48.48%±1.41; inactive+LIRD: 38.79%±0.95; active+LIRD: 47.88±1.40; two-way ANOVA, effect of exercise*LIRD, F_(3,42)_=10.49, p<0.0001) and total vessel length (**Figure 2N**, inactive+dim: 9669μm ±326.20; active+dim: 10472μm ±242.0; inactive+LIRD: 7710μm ±176.20; active+LIRD: 9788μm ±180.30; two-way ANOVA, effect of exercise*LIRD, F_(3,42)_=24.49, p<0.0001); with a significant increase in total number of end points (**Figure 2O**, inactive+dim: 247.20a.u.±11.16; active+dim: 178.9a.u.±9.55; inactive+LIRD: 275.10a.u.±11.31; active+LIRD: 199.40±13.64; two-way ANOVA, effect of exercise*LIRD, F_(3,42)_=13.73, p<0.0001), and lacunarity (**Figure 2P**, inactive+dim: 0.054a.u.±0.0052; active+dim: 0.038a.u.±0.0046; inactive+LIRD: 0.075a.u.±0.0074; active+LIRD: 0.040a.u.±0.0036; two-way ANOVA, effect of exercise*LIRD, F_(3,42)_=8.78, p=0.0001).

### Exercise protects against degradation of neurovascular coupling

Functional hyperemia was performed to quantify changes in venule and arteriole vasodilation in response to photoreceptor stimulation with flickering light (**Figure 3**). The arterial caliber plots across time show that LIRD reduces arterial dilation, an effect prevented in exercised mice (**Figure 3A-E**). With quantification, we found exercise significantly increased arteriole (inactive+dim: 8.18%±1.35; active+dim: 10.39%±1.25; inactive+LIRD: 4.16%±0.65; active+LIRD: 9.34%± 0.68; two-way ANOVA, effect of exercise*LIRD, F_(3,30)_=6.26, p=0.002; **Figure 3E**) and venule diameter percent change in response to flicker stimuli (inactive+dim: 4.44%±0.55 active+dim: 6.53%±0.95; inactive+LIRD: 3.74%±0.57; active+LIRD: 6.52%±0.53; two-way ANOVA, effect of exercise*LIRD, F_(3,32)_=4.14, p=0.014, **Figure 3F**).

**Figure 3.**
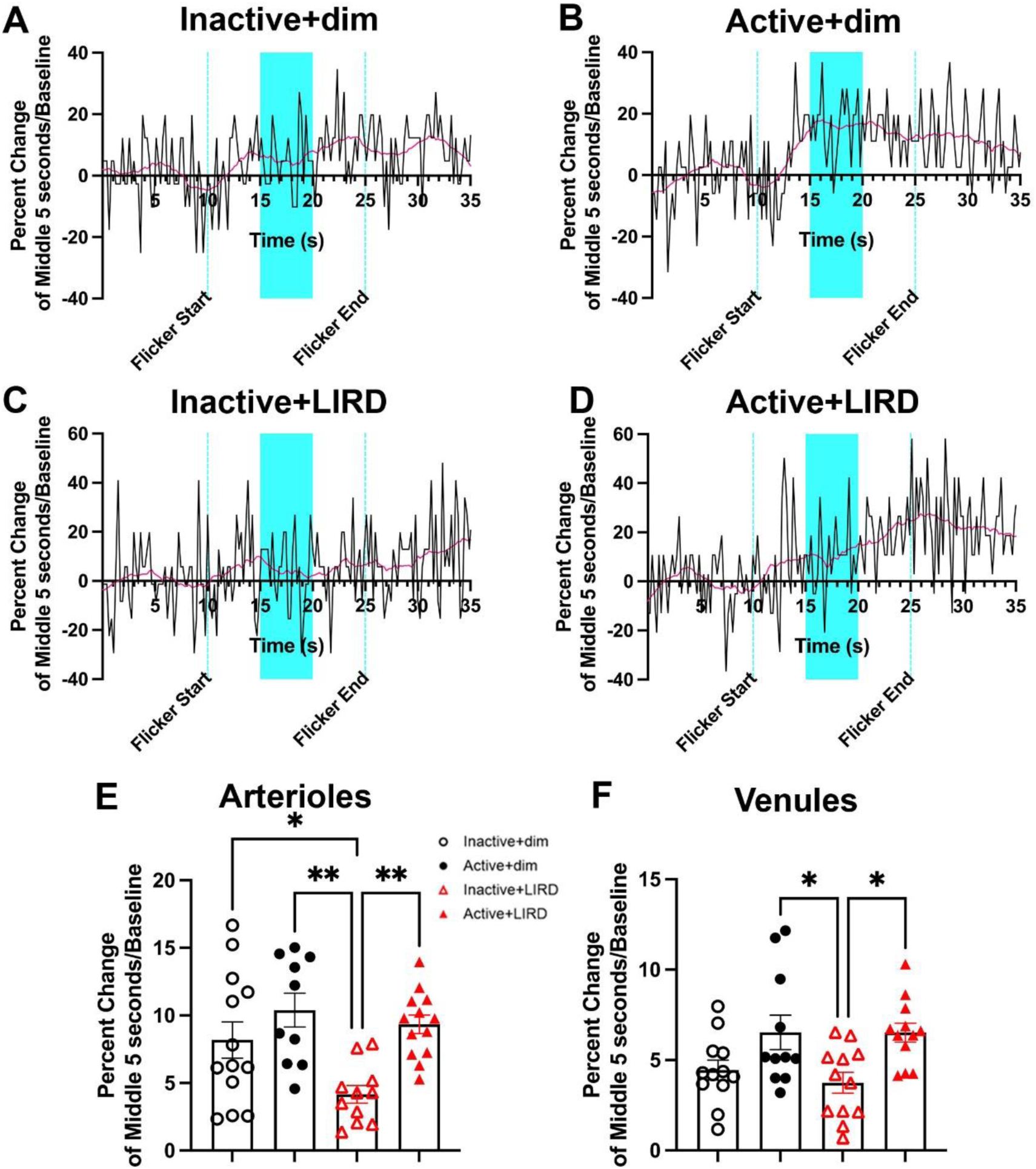
Exercise protects against degradation of neurovascular coupling. Percent change of first order retinal arteriole and venule dilation was measured utilizing flicker induced functional hyperemia. **A-D**, Representative plots of percent change in arteriole caliber over time, with blue highlighting the quantified region. Active+dim and active+LIRD mice had a significant increase in arteriole (**E**) and venule (**F**) dilation compared to inactive groups, with inactive+LIRD arteriole dilation being significantly decreased compared to all groups. Two-way ANOVA with Tukey’s multiple comparison analyses were performed, n=10-13,*p<0.05, **p<0.01.

### Treadmill exercise modestly alters retinal endothelial cell gene expression in active LIRD mice comparable to controls

Isolated retinal CD31+ endothelial cells were probed for genes associated with angiogenesis and neuroprotection: vascular endothelial growth factor receptor 1 (VEGFR1), VEGFR2, vascular cell adhesion molecule 1 (VCAM1), endothelin 1 (EDN1), nitric oxide synthase 3 (NOS3), brain derived neurotrophic factor (BDNF), nuclear factor kappa B subunit 1 (NFKB1), and nuclear factor kappa B subunit 2 (NFKB2). Significant differences were found primarily between active+dim and inactive+LIRD retinas, with active+LIRD retinas largely did not exhibit significant expression changes compared to dim groups; Inactive+LIRD endothelial cells had significantly increased expression of VEGFR1 (**Figure 4A**, two-way ANOVA, effect of exercise*LIRD, F_(3,9)_=3.70, p=0.037) and VCAM1 (**Figure 4C**, two-way ANOVA, effect of exercise*LIRD, F_(3,9)_=5.65, p=0.019). Active+LIRD animals had a significant increased expression of EDN1 compared to inactive+dim (**Figure 4D**, two-way ANOVA, effect of exercise*LIRD, F_(3,9)_=4.26, *p=0.039). Inactive+LIRD isolated endothelial cells also had increased expression of Nfkb1 compared to both dim groups and active+LIRD retina were significantly increased compared to active+dim retina (**Figure 4G**, two-way ANOVA, effect of exercise*LIRD, F_(3,9)_=7.64, *p=0.029, **p=0.0061). No significant differences between groups were found for VEGFR2 (**Figure 4B**), NOS3 (**Figure 4E**), BDNF (**Figure 4F**), NFKB2 (**Figure 4H**).

**Figure 4.**
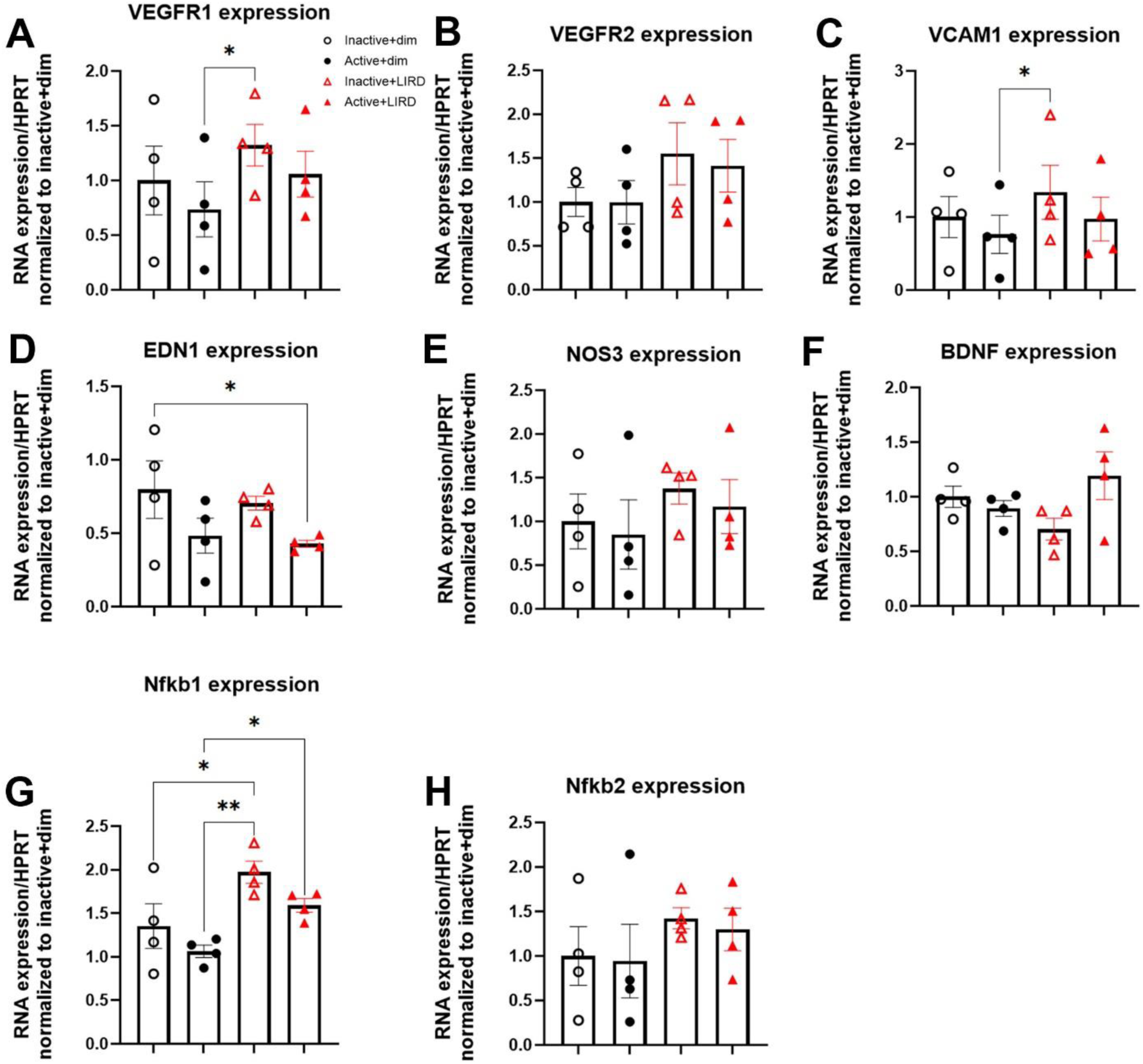
Treadmill exercise modestly alters retinal endothelial cell gene expression in active LIRD mice comparable to controls. Isolated retinal endothelial cells from all experimental groups were probed for genes associated angiogenesis, neuroprotection and inflammation. Inactive+LIRD endothelial cells had significantly increased expression of vascular endothelial growth factor receptor 1 (VEGFR1, **A**) and vascular cell adhesion molecule 1 (VCAM1, **C**) expression compared to active+dim groups, whereas no significance was found between inactive+dim and active+LIRD groups. Inactive+LIRD endothelial cells also had significantly increased expression of Nuclear Factor Kappa B Subunit 1 (Nfkb1, **G**) compared to all groups, additionally active+dim animals were significantly decreased compared to active+LIRD groups. Active+LIRD animals had a significantly decreased expression of Endothelin 1 (EDN1, **D**) compared to inactive+dim groups. No statistical differences were observed in VEGFR2 (**B**), Nitric oxide synthase 3 (NOS3, **E**), and Nuclear Factor Kappa B Subunit 2 (Nfkb2, **H**) expression. Two-way ANOVA with Tukey’s multiple comparison analyses were performed, graphs represent mean values with each data point representing n=5 animals, 10 retinas, n=20 per group, *p<0.05, **p<0.01.

### Active+LIRD mice exhibit a specific circulating cytokine and chemokine profile

The multiplex assay revealed distinct serum protein signatures in a subset of cytokines and chemokines in active mice undergoing retinal degeneration (**Figure 5A-M**). Active+LIRD mice had a significant increase in serum levels of erythropoietin (EPO; **Figure 5B**, two-way ANOVA, effect of exercise*LIRD, F_(3,26)_=3.15,p=0.042), interleukin-1Beta (IL-1β; **Figure 5C**, two-way ANOVA, effect of exercise*LIRD, F_(3,26)_=5.84,p=0.0034), interleukin-6 (IL-6; **Figure 5D**, two-way ANOVA, effect of exercise*LIRD, F_(3,21)_=4.52, p=0.014), chemokine (C-C motif) ligand 12 (CCL12; **Figure 5E**, two-way ANOVA, effect of exercise*LIRD, F_(3,26)_=7.35,p=0.0010), B-lymphocyte chemoattractant (BCA-1/BLC; **Figure 5F**, two-way ANOVA, effect of exercise*LIRD, F_(3,24)_=8.15,p=0.0006), interleukin-21 (IL-21; **Figure 5G**, two-way ANOVA, effect of exercise*LIRD, F_(3,22)_=9.23,p=0.0004), vascular endothelial growth factor-A (VEGF-A; **Figure 5H**, two-way ANOVA, effect of exercise*LIRD, F_(3,25)_=9.47,p=0.0002), and keratinocyte-derived chemokine (KC; **Figure 5N**, two-way ANOVA, effect of exercise*LIRD, F_(3,25)_=10.73,p=0.0001). Both active groups had significant decreased expression in the following cytokines compared to inactive groups: Tumor necrosis factor alpha (TNF-a; **Figure 5I**, two-way ANOVA, effect of exercise*LIRD, F_(3,24)_=6.19,p=0.0029), macrophage inflammatory protein-1 beta (MIP-1β; **Figure 5J**, two-way ANOVA, effect of exercise*LIRD, F_(3,24)_=3.68,p=0.026), interleukin-17C (IL-17C, **Figure 5K**, two-way ANOVA, effect of exercise*LIRD, F_(3,26)_=3.69,p=0.025), tumor necrosis factor receptor superfamily member 5 (CD40/TNFRSF5; **Figure 5L**, two-way ANOVA, effect of exercise*LIRD, F_(3,24)_=6.56,p=0.0021), RANTES (**Figure 5M**, two-way ANOVA, effect of exercise*LIRD, F_(3,26)_=5.096,p=0.0066), interleukin-16 (IL-16, **Figure 5O**, two-way ANOVA, effect of exercise*LIRD, F_(3,26)_=6.82,p=0.0015), eotaxin (**Figure 5P**, two-way ANOVA, effect of exercise*LIRD, F_(3,25)_=8.12,p=0.0006). Matric metalloproteinase-9 (MMP-9; **Figure 5Q**, two-way ANOVA, effect of exercise*LIRD, F_(3,22)_=7.99,p=0.0009) was significantly increased in inactive+LIRD animals compared to all other groups.

**Figure 5.**
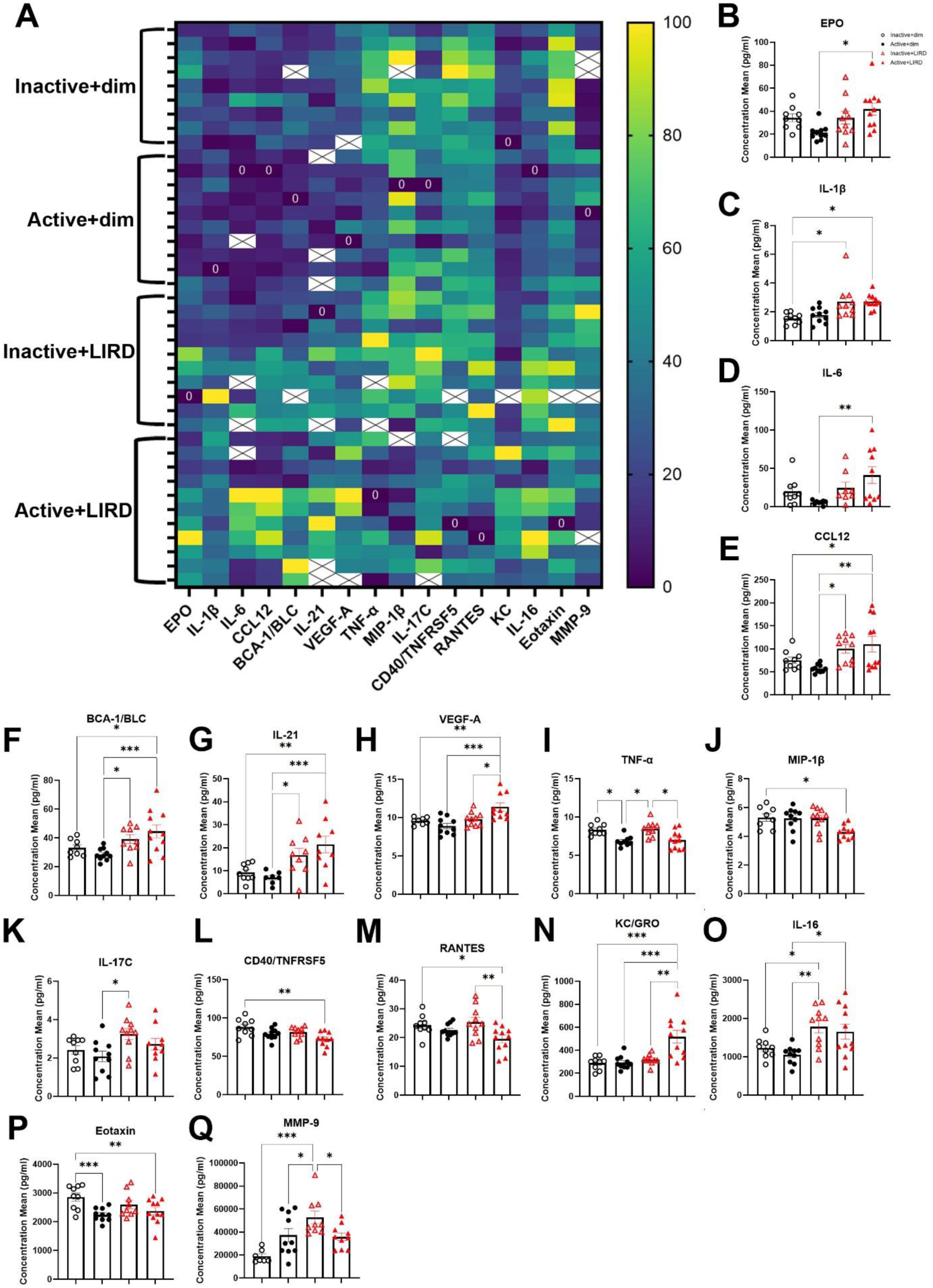
Active+LIRD mice reveal a specific circulating cytokine and chemokine profile. Multiplex analyses quantifying cytokine and chemokine expression related to circulating immune response. Active+LIRD animals have a specific circulating immune response profile compared to all other experimental groups, which support retinal neuron and vascular survival and function during retinal degeneration. This includes chemokines and cytokines involved in macrophage and neutrophil recruitment. Two-way ANOVA with Tukey’s multiple comparison analyses were performed, graphs represent mean values with each data point representing n=9-11 per group, *p<0.05, **p<0.01, ***p<0.001.

## Discussion

To date, there are minimal studies investigating the neuroprotective effects of exercise on retinal vasculature during RD^12^. Our results demonstrate that treadmill exercise impacts retinal vascular morphology, function, endothelial gene expression, and the expression profile of circulating factors in the LIRD model of photoreceptor degeneration.

To assess *in vivo* retinal vascular plexi structure, we used OCT-A to segregate and quantify vascular plexi morphology. Inactive+LIRD mice showed breakdown of the vascular beds that support the inner retina, quantified by a significant decrease in vessel density, total vessel length, and an increase in the total number of end points as well as lacunarity (a measurement of areas lacking vasculature) compared to all other groups. Collectively, these data indicate that vascular remodeling and degradation occurs with photoreceptor degeneration and that treadmill exercise prevents vascular breakdown with active+LIRD retinas being statistically similar to active+dim groups.

Retinal flicker-evoked vasodilation, or retinal functional hyperemia, is a powerful non-invasive tool to detect vascular dysregulation^15,16^. A reduction in retinal functional hyperemia is one of the earliest retinal functional changes observed in patients with RD that primarily target retinal vasculature such as age-related macular degeneration and diabetic retinopathy^17–19^. Our results demonstrate retinal arteriole vasodilation was significantly decreased in inactive+LIRD mice compared to all other groups, and retinal venule vasodilation was significantly decreased compared to active groups. Active+LIRD animals were statistically indistinguishable from dim groups. Importantly, our findings of significantly decreased vasodilation in inactive+LIRD mice aligns with previous results described in patients with RD and supports retinal vascular functional losses occur simultaneously with photoreceptor degeneration^2,6^. These data suggest that treadmill exercise maintains retinal vascular function and improves neurovascular coupling during retinal degeneration.

To investigate potential gene expression changes associated with angiogenesis and neuroprotection, retinal endothelial cells were isolated from whole retina. Vascular endothelial growth factors (VEGFs) are the principal drivers of angiogenesis through the binding of their tyrosine kinase receptor (VEGFRs) to provoke various downstream effects in endothelial cells^20^. Isolated retinal endothelial cells from inactive+LIRD animals revealed significantly increased expression of VEGFR1 compared to active+dim animals. Although not statistically significant, inactive+LIRD isolated retinal endothelial cells also trended towards having increased expression of VEGFR2, which is the main receptor involved in the progression of angiogenesis. VEGFR1 is known as the decoy receptor for VEGF, preventing it from interacting with VEGFR2^20^. VCAM1 was significantly increased in inactive+LIRD retinas compared to active+dim retinas, which supports immune cell-blood vessel interaction. Recently, increased VCAM1 expression was found to potentially contribute to the development of macular fibrosis in neovascular age-related macular degeneration patients^21^. EDN1, which regulates blood vessel dilation and constriction, in endothelial cells from active+LIRD was significantly decreased compared to inactive+dim animals. Nfkb1, a regulator of pro-inflammatory cytokines, was also significantly altered across groups, with endothelial cells from inactive+LIRD having significantly increased expression^19^. Additionally, active+LIRD animals had a trend in increased expression of BDNF, although not significant.

The injury expression profile in inactive+LIRD animals had a significant increase in IL-1β compared to inactive+dim animals, a significant increase in CCL12, BCA-1/BLC, IL-21, IL-17C expression compared to active+dim groups, and a significant increase in MMP-9 compared to all groups. IL-16 was increased in both LIRD groups. Active+LIRD animals presented with a specific circulating chemokine and cytokine expression profile in serum, that was distinct from all other experimental groups. Active+LIRD animals had a significant increase in: EPO, IL-1β, IL-6, CCL12, BCA-1, IL-21, VEGF-A, KC; and a decrease in: TNF-α, MIP-1β, CD40, and eotaxin, compared to all other groups. This specific combination of upregulated and downregulated circulating factors may give insight for a distinct inflammatory response profile that affords and supports retinal neuroprotection induced by exercise, through the recruitment and engagement of specific immune cells such as microglia, macrophages, and neutrophils.

Collectively, this work suggests that exercise-induced retinal neuroprotection is accompanied by preservation of retinal vasculature. We have shown that animals undergoing retinal degeneration benefit from exercise to help maintain vascular morphology, improve arteriole and venule vasodilation, and modestly alter retinal endothelial gene expression towards a neuroprotective profile. Our work also suggests that exercise induces a distinct immune response profile, with specific chemokines and cytokines being up and downregulated. Overall, these data reveal some of the structural, functional, and molecular mechanisms involved in exercise-induced retinal neuroprotection as it pertains to retinal vasculature in a retinal degeneration model.

